# The first insight into the genetic structure of the population of modern Serbia

**DOI:** 10.1101/2020.12.18.423408

**Authors:** Tamara Drljaca, Branka Zukic, Vladimir Kovacevic, Branislava Gemovic, Kristel Klaassen-Ljubicic, Vladimir Perovic, Mladen Lazarevic, Sonja Pavlovic, Nevena Veljkovic

**Author notes:** Joint corresponding authors.

## Abstract

The complete understanding of the genomic contribution to complex traits, diseases, and response to treatments, as well as genomic medicine application to the well-being of all humans will be achieved through the global variome that encompasses fine-scale genetic diversity. Despite significant efforts in recent years, uneven representation still characterizes genomic resources and among the underrepresented European populations are the Western Balkans including the Serbian population. Our research addresses this gap and presents the first ever dataset of variants in clinically relevant genes in the population sample of contemporary Serbia. A few variants significantly more frequent in the analyzed sample population compared to the European population as a whole are distinguished as its unique genetic determinants. We explored thoroughly their potential functional impact and its correlation with the health burden of the population of Serbia. Our variant’s catalogue improves the understanding of genetics of modern Serbia, contributes to application of precision medicine and health equity. In addition, this resource may also be applicable in neighboring regions and in worldwide functional analyses of genetic variants in individuals of European descent.

## Introduction

In the era of high-throughput next generation sequencing (NGS) technology, the common goal for clinical use of sequencing data is the identification of pathogenic variants that can affect an individual’s health through linking genes and diseases [1–3]. The importance of elucidating fine-scale genetic diversity lies in understanding the genomic contribution to certain conditions, response to treatments, and in the application of acquired knowledge to clinical care and well-being of all people [4]. In the health context, equity will be achieved through the unbiased implementation of genomic medicine and evenly balanced structure of genomic analyses that comprise human specificities [5]. The misclassification of variants coming from data that do not include diverse subpopulations can potentially lead to misinterpretation of causative factors of disease and inadequate treatments of individuals from underrepresented segments [6]. Although large-scale variome studies, such as the 1000 Genomes Project [7], Exome Aggregation Consortium (ExAC) [8] and Genome Aggregation Database (gnomAD) [9] have widely expanded our horizons on human diversity, numerous studies [10–13] are demonstrating that there are many more population-specific variations than have been captured through these initiatives. In recent years, significant effort has been invested in addressing gaps in the composition of the global genomic landscape [4].

Many countries have performed national studies to create population-specific variant panels and supplement the universal reference genome. These studies have a common aim in understanding genetic variability at the population level, as well as understanding and interpreting pathogenic variants and prioritizing candidate disease-causing genetic variation. The Genome of the Netherlands [14] and the Danish Reference Genome Project [15] were among the first European projects for creating a national pan-genome. Both projects were based on trios and aimed at applying a newly acquired dataset as a reference for clinical and medical-sequencing projects. SweGen [16] and Iceland’s project [17] are other examples of national projects intended to assess the genetic variability at a more detailed level in order to establish a control dataset for the local population. Another national project, the UK Biobank [18,19] is the most ambitious contemporary population genetics study set up to potentiate genetic and non-genetic determinants of the disease.

Although European population genetics has been largely studied and well described, the Western Balkans including Serbia are underrepresented in the majority of cohorts. Modern Serbia is a landlocked Western Balkan country in south eastern Europe. Its population of about 7 million citizens is demographically old [20] with a health profile predominantly burdened with cardiovascular diseases and cancer [21]. However, thus far, there have been no reports on the contemporary Serbian variome. Most of the research on Balkan populations was conducted on uniparentally inherited markers such as mitochondrial DNA (mtDNA) and the Y chromosome, focusing on population descent and haplogroup diversity [22–24]. One recent exception is a report that describes the sequencing and analysis of a genome from a contemporary individual of Serbian origin and which introduces tens of thousands of previously unknown variants [25].

In this work, our focus was on common variants and their functional impact in the Serbian population sample. We created a catalogue of variants called after the clinical exome sequencing of 147 individuals from Serbia. Unique genetic characteristics of the studied population were identified as variants that are frequent in the Serbian sample but much less frequent in the European population. Our data set not only provides the first reference point for genomic studies and precise medicine in Serbia, but also supports worldwide functional analyses of genetic variants in individuals of European descent.

## Results and Discussion

### Description and functional prediction of variants in the Serbian population sample

After variant calling from a dataset that included sequencing data of 147 individuals, followed by sample and variant QC filtration, we obtained a final multisample set reduced to 144 data samples. The final set has 47,324 elements, of which SNVs represent 95.80% (Fig. 1.a). The transition-transversion ratio (Ti/Tv), as the measure of the overall SNV quality, was evaluated before and after QC and substitution type distribution as shown in (Fig. 1.b). The Ti/Tv ratio increased from 2.92 to 2.97 which is in accordance with the quality expected for the exome data.

**Figure 1.**
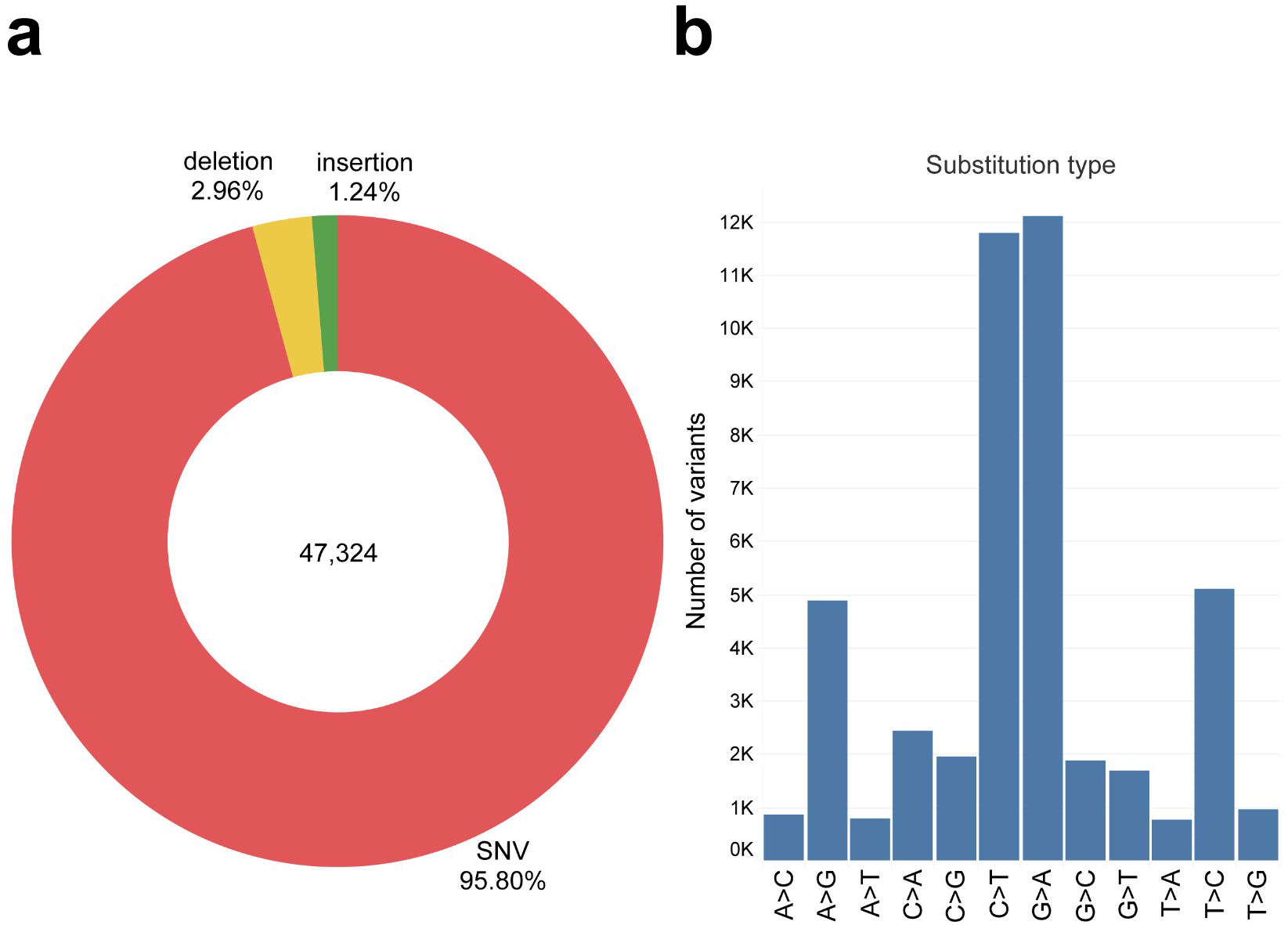
Serbian population sample variant classes and types. **a.** Variant distribution by class shows that the vast majority of the total number of variants (47,324) are SNVs, followed by a significantly lower percent of deletions and insertions **b.** SNV substitution type distribution

Functional classification showed that variants with moderate functional impact are the most abundant (19,918), whereas high impact variants make up 2.31% (1,093) (Fig. 2.a, Supplementary Table 1). Each functional impact category was divided by minor allele frequency (MAF) and singleton and doubleton subcategories (Fig. 2.b). In our dataset 11.94% of the total number of variants was rare (MAF≤1%), while the number of common variants (MAF≥5%) on which we focused our research was 13,362 (28.24%). We calculated the number of singletons (Major Allele Count, MAC = 1) and private doubletons (MAC = 2) separately and singletons make up 39.26% of the total number of variants, while private doubletons make up 1.03%. The cause of the high percent of singletons and private doubletons may be attributed to the small overall sample size [26].

**Figure 2.**
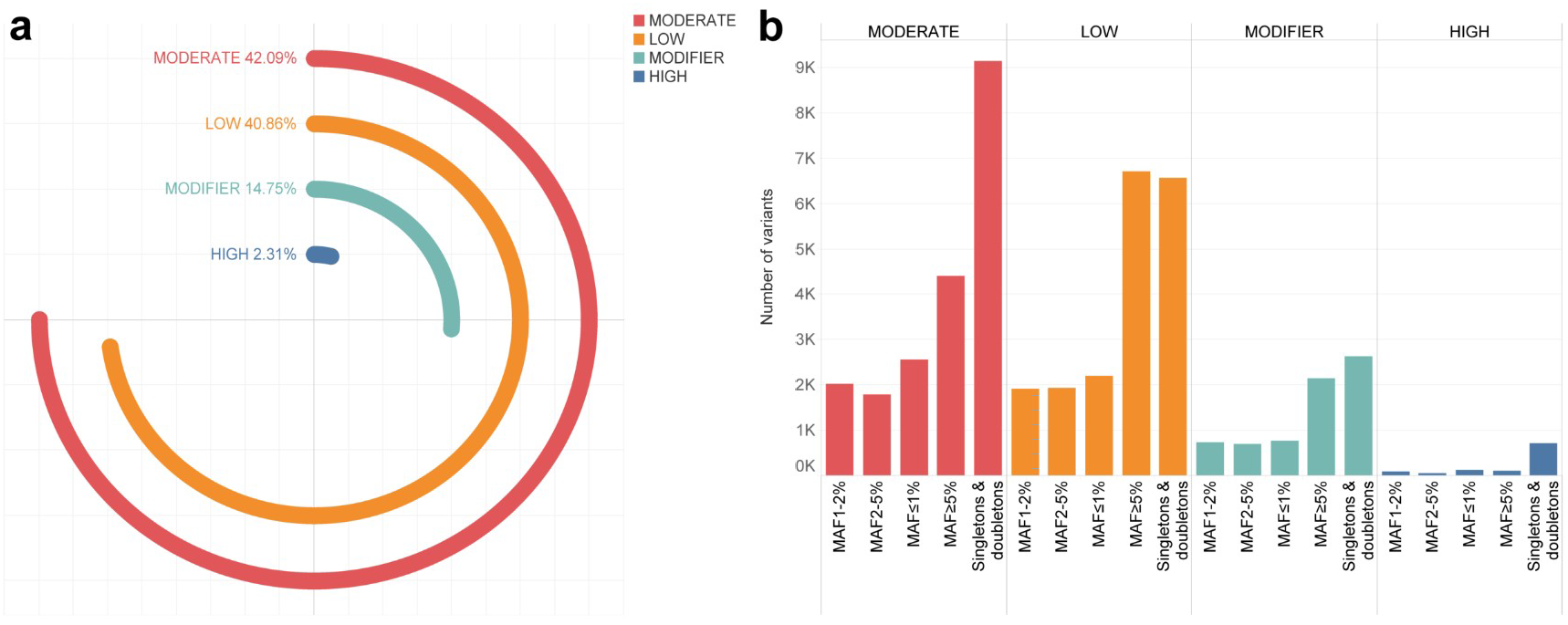
Functional effects of variants in the Serbian population sample. **a.** The distribution of variants across predicted categories. Modifiers and high functional impact variants are predicted for 17.06 % of variants. The vast majority of variants are predicted to have moderate and low functional impact. **b.** Functional impact categories divided into MAF subcategories. Value on the y-axis represents the number of variants in each MAF subcategory represented on the x-axis are: MAF ≤1%, MAF1-2%, MAF2-5%, MAF ≥5% and singletons (MAC=1) and private doubletons (MAC=2) subcategory.

Functional annotation using the Ensembl Variant Effect Predictor (VEP) [27] included embedded pathogenicity predictions with SIFT v5.2.2 [28] and PolyPhen-2 v2.2.2 [29]. Pathogenicity prediction using SIFT was obtained for 40.33% (19,089) of the total number of SNVs and for 40.53% (19,183) using the PolyPhen-2 tool (Supplementary Fig. S3). The majority of variants are classified in similar categories by both tools (Fig. 3), i.e. variants predicted to be tolerated by SIFT were also seen as benign by PolyPhen-2 (49.29%), whereas most of those predicted to be deleterious by SIFT are perceived as damaging or probably damaging by PolyPhen-2. However, 9.42% of variants are cross-classified as deleterious and benign, while 3.25% variants are both tolerated and probably damaging (Supplementary Table S2). This SNVs set might be of particular interest because it is possible that their functional effects are subtle and thus, their significance may remain unexplored. In our future research we should explore these variants, particularly the mixed classified common ones (MAF≥5%).

**Figure 3.**
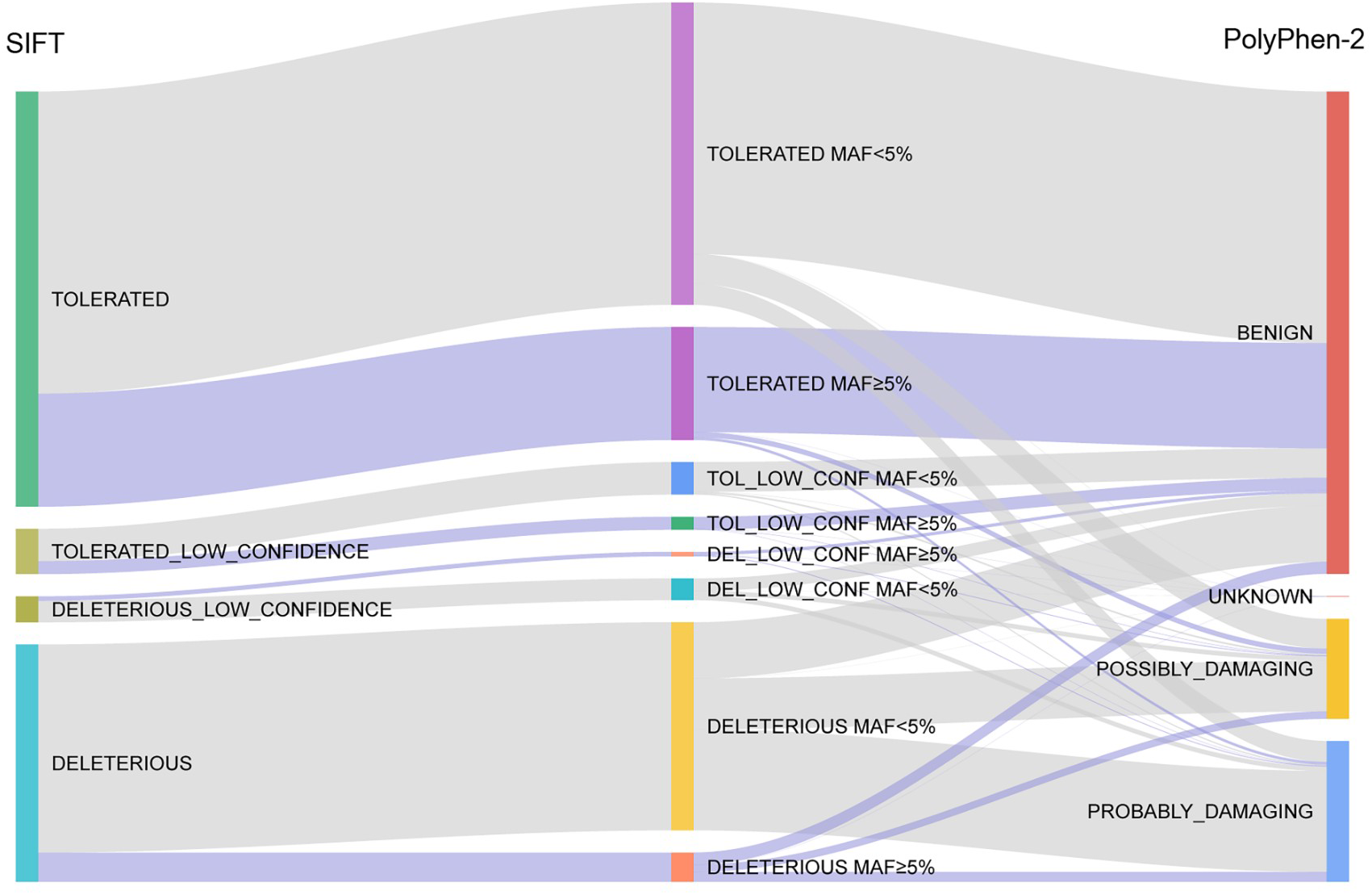
SIFT and PolyPhen-2 functional impact and different predictions. The figure presents the flow between the SIFT tool [28] (left side of the chart) and PolyPhen-2 tool [29] (right side of the chart) predictions. SNVs are differentiated between common variants, MAF≥5% (grey lines) and rare variants, MAF<5% (violet lines).

### Ancestry analysis

The Balkan Peninsula, encompassing the larger part of Serbian territory is situated at the crossroads of Central and Southeast Europe and was one of key areas in major migratory events that occurred after the Last Glacial Maximum [30–32]. The area of present-day Serbia has been inhabited since the Paleolithic Age [33]. As for the Neolithic Age, analysis of genome-wide DNA polymorphisms of populations bordering the Mediterranean coast, including Serbia, confirmed the hypothesis that the maritime coastal route was mainly used for the migration of Neolithic farmers to Europe [34]. Here, we investigated the ancestry pattern in this study sample by combining pruned and common SNVs (MAF≥5%) of the Serbian population sample with SNVs of 1kGP as a reference dataset. Principal component analysis (PCA) revealed that the Serbian cluster overlaps with the European sample that clearly differentiates from the other super populations of 1kGP (Fig. 4).

**Figure 4.**
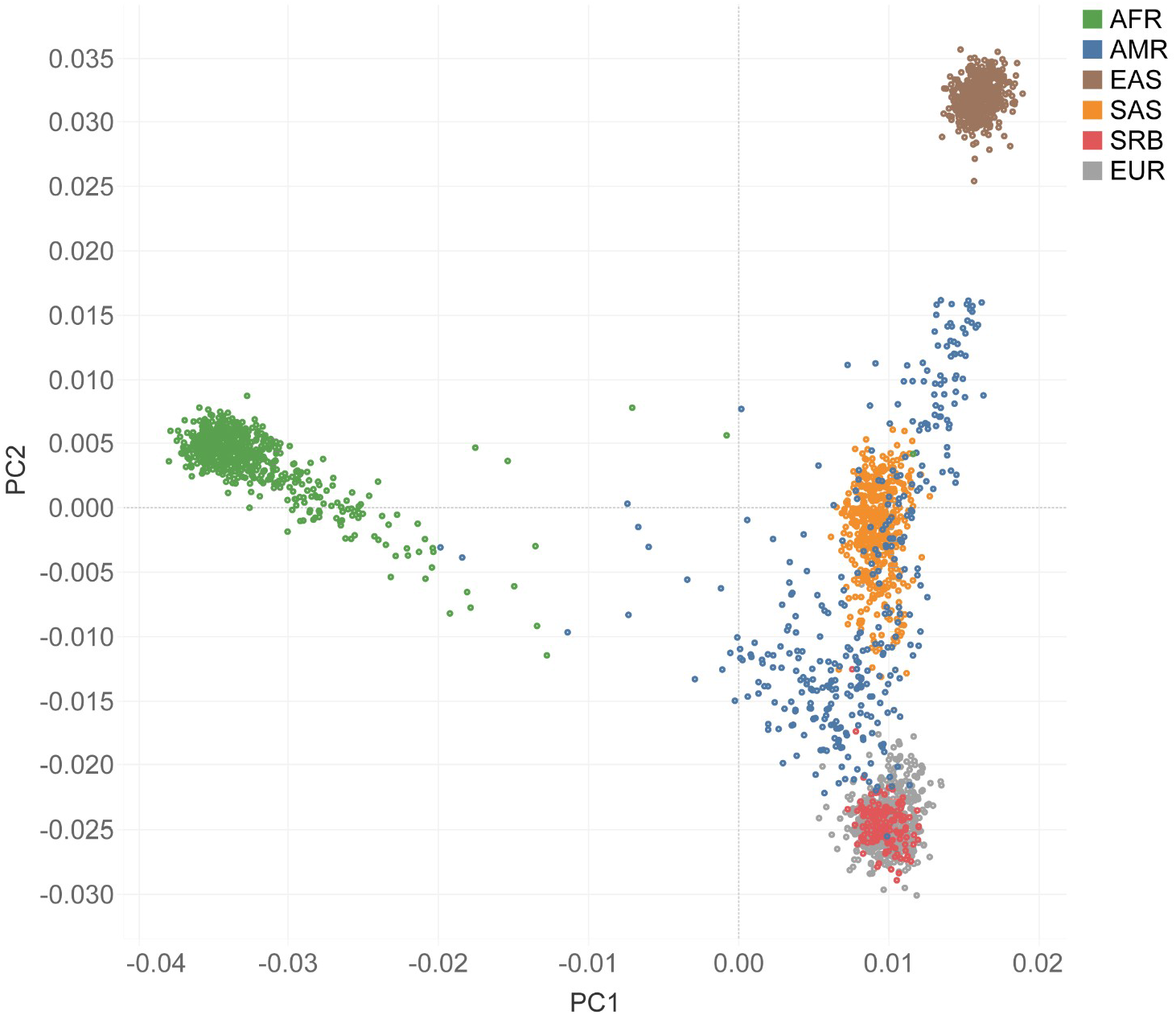
Principal component analysis (PCA) of the Serbian population sample. PCA of the study data combined with the 1000 Genomes Project Phase 3 data (AFR = African, AMR = Admixed American, EAS = East Asian, SAS = South Asian, EUR = European). Serbian population sample data, labeled as Serbian (SRB), overlaps with the European (EUR) ancestry cluster.

### Distribution of novel variants in the Serbian population sample

Our final dataset was intersected with reference databases 1kGP Phase 3, European population [7], gnomAD v3.0 [9], and NHLBI ESP (https://evs.gs.washington.edu/EVS/) and revealed that 4972 (10.5%) variants are not present in any of them. These variants are referred to as novel and their presence in different databases is shown in Fig. 5. As expected, our dataset overlaps the most with gnomAD’s dataset which is the largest and which was mapped to the hg38 reference genome.

**Figure 5.**
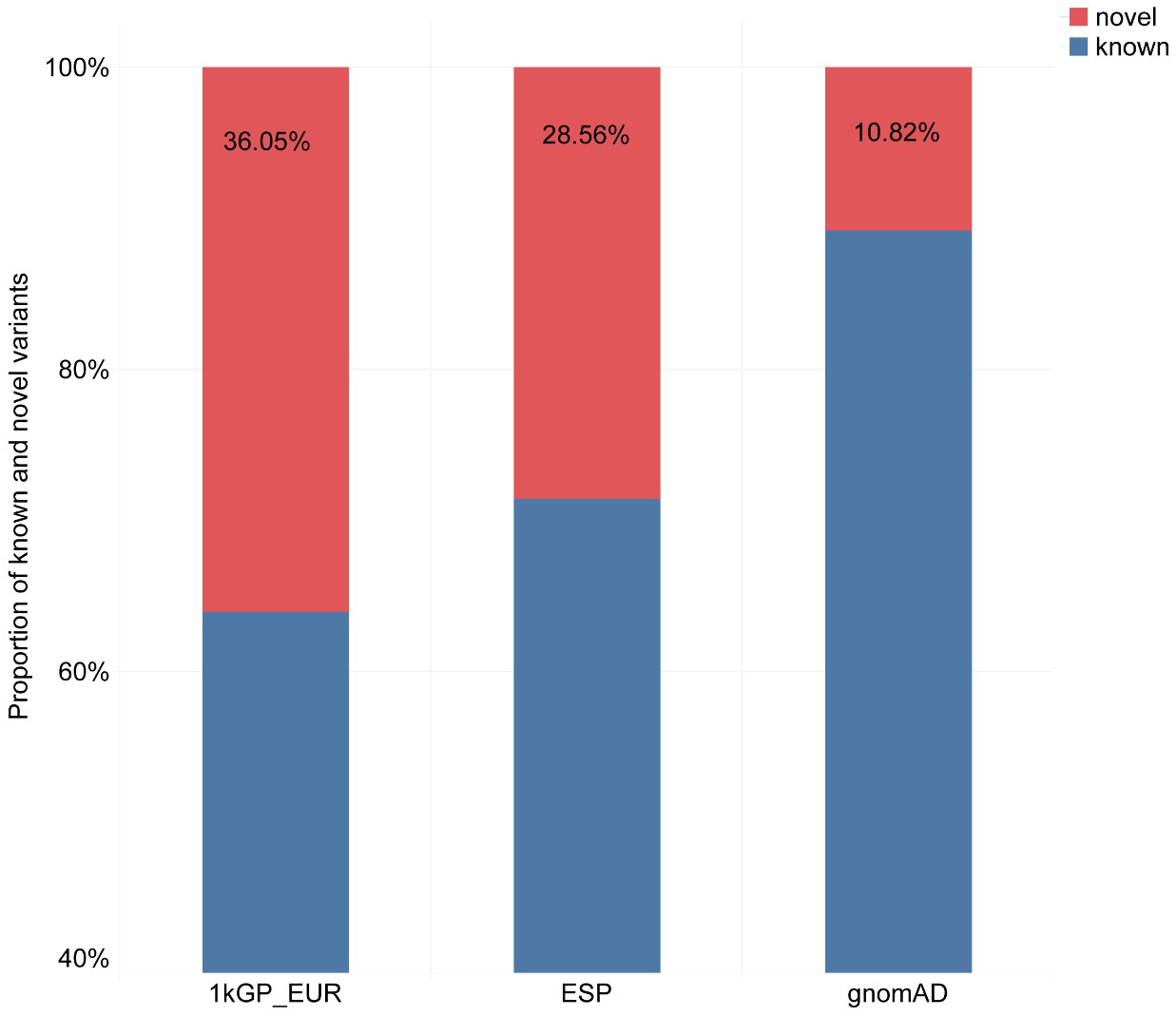
Ratio of known and novel variants in the Serbian population sample per reference database: 1kGP_EUR, gnomAD and NHLBI ESP.

Analyses of functional effects of novel variants revealed that the majority of novel variants (450) are in the category of HIGH impact variants, followed by the MODIFIER category. Furthermore, considering allele frequency and allele count, the majority of novel variants are rare in the Serbian population sample (Fig. 6).

**Figure 6.**
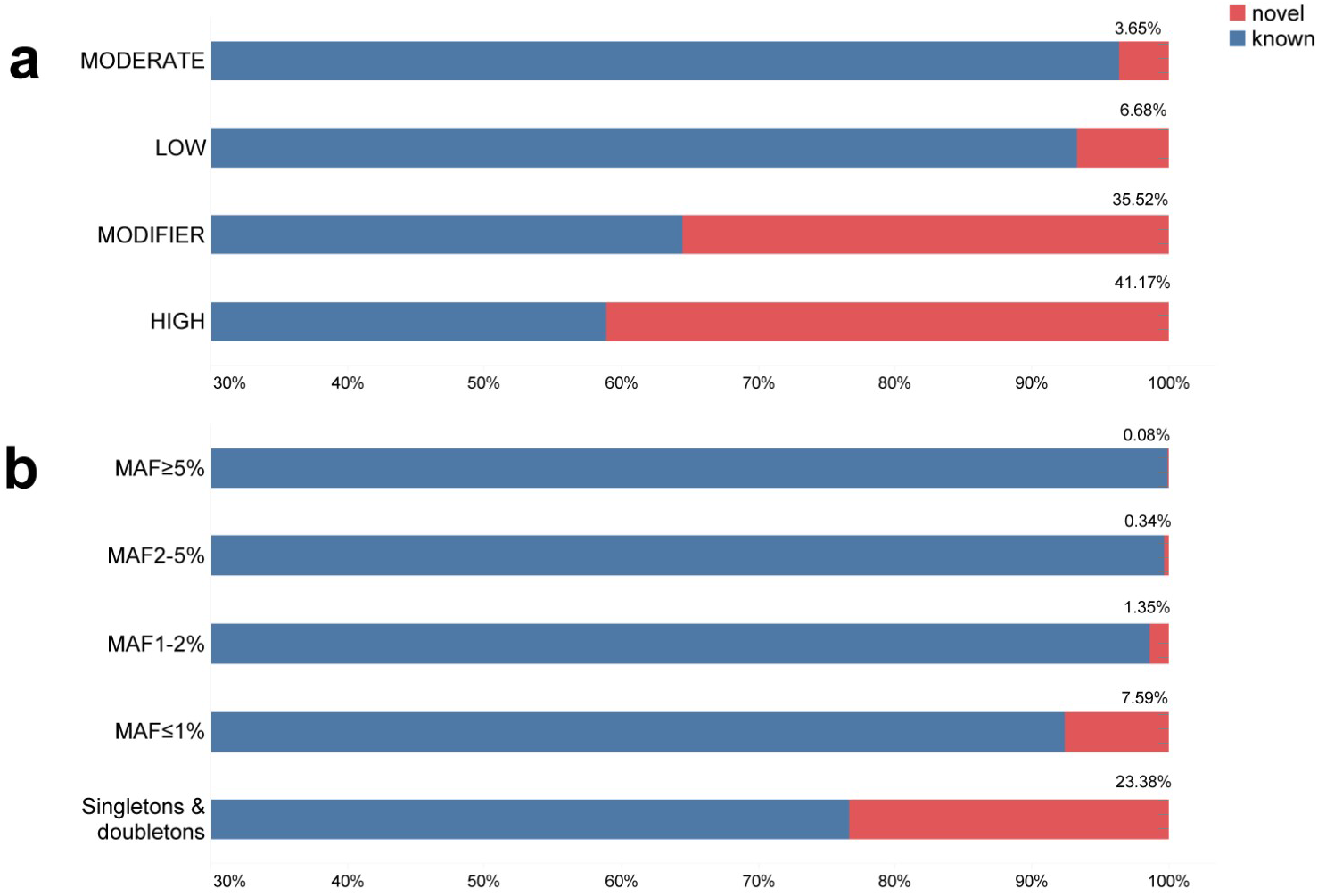
Novel variants in the Serbian population sample classified by predicted functional impact. **a.** Percent of novel variants by functional impact relative to known variants found in all databases after intersection. **b.** Percent of novel variants distributed across allele frequency categories.

### Missense variants frequent in the Serbian population sample: Case studies

Using the CNVkit 0.9.1 toolkit [35] we were able to determine the sex distribution in the Serbian sample. After the QC filtration, we excluded three samples due to Het/Hom deviation and we kept 61 female sample out of 62, and 83 male samples out of 85. Furthermore, we analysed the distribution of common variants found in Serbian sample in the remaining female and male samples (Table 1, Supplementary Table S3). As the literature search of overrepresented variants revealed that they were almost not investigated at all, we explored annotations of genes that harbor these variants in the Gene Ontology database in order to better understand the processes and pathways that might be affected. In these analyses we were restricted to the sub-ontology biological processes (GO-BPO) and found that five genes are involved in the immune response and two genes participate in chemical synaptic transmission (Supplementary Table S5). Four genes, MYO15A, RHPN2, BTNL2 and GAL3ST3, do not have annotations in GO-BPO. One interesting coincidence is that the protein product affected by the variant that distinguishes the most the studied population from other Europeans has the same name as an individual from the largest ethnic group in Serbia, the Serbs. The PSPH protein product SERB is a phosphoserine phosphatase and a member of the haloacid dehalogenase superfamily of hydrolytic dehalogenases [36].

**Table 1.** Top 5 variants with the highest fold increase in the Serbian population sample compared to Europeans and their sex representation. Variant MYO15A (position 17:181208) does not have rsID in dbSNP138.

| Gene | Variant rsID<br>dbSNP138 | Substitution<br>type | % of<br>variants in<br>the female<br>sample | % of<br>variants in<br>the male<br>sample | Fold<br>increase |
| --- | --- | --- | --- | --- | --- |
| PSPH | rs79451216 | missense | 36% | 30% | 163 |
| PCDHB4 | rs147934595 | missense | 43% | 46% | 74 |
| MRC1 | rs71497224 | synonymous | 93% | 94% | 70.73 |
| MYO15A | - | synonymous | 0 | 1% | 50 |
| KIR2DL1 | rs79002558 | missense | 15% | 11% | 33 |

Missense variants in these genes were further analyzed by using MutPred2 [37], a tool that predicts not only the pathogenicity as PolyPhen-2 [29] and SIFT [28] but also the molecular mechanisms underlying the effects of variants predicted to be pathogenic (Table 2, Supplementary Table S4).

In the subsequent paragraphs we review a few genes that harbor variants characteristic of the population of Serbia.

The PSPH (Phosphoserine Phosphatase) gene codes for a member of the SerB protein family, a phosphoserine phosphatase involved in the biosynthesis of serine [38]. A recent study by Jia et al. [39] showed that the PSPH loci is associated with the glycine level, while Byers et al. [40] reported a decreased glycine level in a patient with PSPH mutations (V44G and G141S). A variant rs79451216 in PSPH, identified as frequent in the Serbian population sample, encompasses two alleles leading to AAS of arginine at position 49 in the protein sequence, R49W and R49G. Of note, this variant is present in 22 out of 61 female samples and in 25 out of 83 male samples. However, sex differences for this and other variants have to be further investigated. MutPred2 showed that these substitutions affect the PSPH protein functions, while molecular mechanisms underlying this disturbance were predicted to be associated with phosphorylation and cleavage of the PSPH protein (Table 2). Thus far, there is no information about the effect of these variants at the level of metabolites affected by PSPH, but their proximity to the already described glycine decreasing variant [40] can lead to the assumption of the same effect. Since glycine was shown to have antihypertensive and atheroprotective properties, as well as, to reduce risk of acute myocardial infarction [41,42], gene variants lowering the glycine level in blood might increase susceptibility to various cardiovascular diseases. Ischemic heart disease and cerebrovascular diseases are the most dominant causes of death in Serbia, [21,43] and the rate of ischemic heart disease in Serbia was higher in comparison with all other European regions [21]. Although this can be attributed to several factors, our results for the first time implicate variations in the PSPH gene as a possible contributor to the high incidence of cardiovascular diseases in Serbia.

The KIR2DL1 (Killer cell immunoglobulin-like receptor 2DL1) codes for one of the receptors on natural killer cells, inhibiting their activity. KIR genes are located in one of the most variable regions of the human genome. Variability of KIR genes is represented by both the presence/absence of genes and sequence polymorphism, leading to high diversity among individuals, as well as different populations [44]. Ligands for this protein are the Human Leucocyte Antigen C (HLA-C) molecules and this interaction is involved in the immune response and dealing with various pathogen infections, including the Hepatitis C Virus (HCV) [45]. Variants in both KIR and HLA-C genes, as well as interleukin 28B (IL28B), affect the course of the HCV infection and the response to therapy [46,47]. Genotyping of these genes is therefore highly recommended in the clinical circumstances involving decisions about the anti-HCV treatment [46]. Since a variant in KIR2DL1 was found to have a 33-fold increase in frequency in the Serbian population sample compared to European populations of 1kGP it should be taken into account when genotyping HCV infected patients in Serbia for this gene. Additionally, this gene variant leads to an amino acid change that was shown in the MutPred2 analysis to be of functional importance (Table 2). Therefore, further consequences of this change for the interaction of KIR2DL1 and HLA-C, as well as additional effects on the immune response in the Serbian population should be tested. Of note, this variant was found in 9 out of 61 female samples and in 9 out of 83 male samples.

Finally, we identified two overrepresented variants in BTNL2 and HLA-DQB1 genes that are according to the literature linked to sarcoidosis, a multiorgan granulomatous inflammatory disease primarily affecting the lungs, but also lymph nodes, liver, spleen, skin, eyes, muscles, brain, kidneys and heart [48]. The BTNL2 (Butyrophilin-like 2) gene codes for a member of the immunoglobulin superfamily, which acts as an inhibitor of T cell activation [49]. BTNL2 is a susceptibility and progression factor for pulmonary sarcoidosis [50] and polymorphisms in this gene are associated with phenotype expression of sarcoidosis [50]. Similarly, an HLA-DQB1 haplotype is strongly associated with severe sarcoidosis [51]. MutPred2 analysis showed that two variants in the BTNL2 and HLA-DQB1 genes, leading to three AAS – S334L and S334W in BTNL2 and D89N, affect the functions of these proteins (Table 2). Pulmonary sarcoidosis in Serbian patients is most often followed by ocular sarcoidosis, as the first most common site of extrapulmonary sarcoid manifestations [52]. Some clinical features of these patients differ from those in other European populations, with more common neuro-ophthalmologic lesions [52]. This difference could be associated with variants in BTNL2 and HLA-DQB1, which we found to be frequent in the Serbian population sample and functionally important, and should be further investigated. The HLA-DQB1 rs41552812 variant was found in 13 out of 61 female samples and in 19 out of 83 male samples. There is an additional example of specificity of the Serbian genomic profile related to sarcoidosis [53].

**Table 2.** Variants detected as frequent in the Serbian population sample that MutPred2 [37] predicted as affecting protein function.

| Gene | Variant | AAS | MutPred2 - Molecular mechanisms |
| --- | --- | --- | --- |
| PSPH | rs79451216 | R49W | ● Loss of relative solvent accessibility<br>● Loss of ADP-ribosylation at R49<br>● Altered metal binding |
|  |  | R49G | ● Loss of relative solvent accessibility<br>● Loss of helix<br>● Gain of ADP-ribosylation at R50<br>● Altered metal binding, gain of methylation at R50 |
| KIR2DL1 | rs79002558 | S88R | ● Loss of relative solvent accessibility<br>● Altered trans-membrane protein<br>● Altered ordered interface<br>● Gain of allosteric site at R89<br>● Gain of ADP-ribosylation at S88<br>● Altered DNA binding<br>● Altered metal binding<br>● Loss of N-linked glycosylation at N84 |
| BTNL2 | rs28362679 | S334L | ● Altered transmembrane protein<br>● Loss of B-factor |
|  |  | S334W |  |
| HLA-DQB1 | rs41552812 | D89N | ● Altered ordered<br>● Gain of relative solvent accessibility |

## Conclusion

Large-scale variome studies have significantly increased our understanding of the diversity in the human population, however, its composition is still broadly biased towards some populations.

In this study we aimed to address the gap in the European genomic landscape and to the best of our knowledge provided the first ever dataset of variants in the population of contemporary Serbia. Variants are described in detail according to allele frequency, presence in key population databases and functional impact as interpreted by several state-of-the-art tools. The common SNV dataset generally overlaps with the European cluster, but we also discerned and reported some unique genetic characteristics as several variants that are significantly overrepresented in the Serbian population sample. These insights will be further evaluated and broadened in larger genomic studies linking genes and diseases. Nevertheless, the variant’s catalogue obtained contributes to our understanding of the genetics of modern Serbia and more adequate functional interpretation in the context of precision medicine and health equity.

## Methods

### Study population

The study cohort consists of 147 unrelated individuals from Serbia. This study was approved by the Ethics Committee of the Institute of Molecular Genetics and Genetic Engineering, University of Belgrade, in accordance with the guidelines of the 1975 Declaration of Helsinki (6th revision, 2008). Informed consent was obtained from all participants included in this study.

### DNA Sequencing

Total genomic DNA was isolated from peripheral blood using the QIAamp DNA Blood Mini Kit (Qiagen, Hilden, Germany). Sequencing was performed in a time span from 2014 to 2020. Sequencing technology was MiSeq Illumina using the TruSight-One Illumina (May 2014) sequencing panel for target exome sequencing, which consists of ~4800 disease-associated genes and ~12 Mb genomic content.

### Variant Calling

Germline SNP and Indel variant calling was performed following the Genome Analysis Toolkit (GATK, v4.1.0.0) best practice recommendations [54]. Raw reads were mapped on the UCSC human reference genome hg38 using a Burrows-Wheeler Aligner (BWA-MEM, v0.7.17) [55]. Optical and PCR duplicate marking and sorting was done using Picard (v4.1.0.0) (https://broadinstitute.github.io/picard/). Base quality score recalibration was done with the GATK BaseRecalibrator resulting in a final BAM file for each sample. The reference files used for base quality score recalibration were dbSNP138, Mills and 1000 genome gold standard indels and 1000 genome phase 1, provided from the GATK Resource Bundle (last modified 8/22/16).

After data pre-processing, variant calling was done with the Haplotype Caller (v4.1.0.0) [56] in the ERC GVCF mode to generate an intermediate gVCF file for each sample, which were then consolidated with the GenomicsDBImport (https://github.com/Intel-HLS/GenomicsDB) tool to produce a single file for joint calling. Joint calling was performed on the whole cohort of 147 samples using the GenotypeGVCF GATK4 to create a single multisample VCF file.

Considering that target exome sequencing data in this study does not support Variant Quality Score Recalibration, we selected hard filtering instead of VQSR. We applied hard filter thresholds recommended by GATK to increase the number of true positive and decrease the number of false positive variants.

All the previous steps were coordinated using the Cancer Genome Cloud Seven Bridges platform [57].

### Quality Control and Annotation

To assess the quality of the obtained set of variants, we calculated per-sample metrics with Bcftools v1.9 (https://github.com/samtools/bcftools), such as the total number of variants, mean transition to transversion ratio (Ti/Tv) and average coverage per site with SAMtools v1.3 [58] calculated for each BAM file. We calculated the number of singletons and the ratio of heterozygous to non-reference homozygous sites (Het/Hom) in order to filter out low-quality samples. Samples with the Het/Hom ratio deviation were removed using PLINK v1.9 (www.cog-genomics.org/plink/1.9/) [59]. We marked the sites with depth (DP) < 20 and genotype quality (GQ) <= 20 and excluded variants where more than 70% of genotypes did not pass the filters.

Deviation from the Hardy-Weinberg equilibrium (HWE) was calculated using VCFtools v0.1.13 [60] with a threshold for HWE ofp < 0.0001 below which the variants were excluded. All variants failing the quality control (QC) steps were removed. All subsequent analyses were performed using clean post-QC datasets.

We used the Ensembl Variant Effect Predictor (VEP, ensembl-vep 90.5) [27] for functional annotation of the final set of variants. Databases that were used within VEP were 1kGP Phase3, COSMIC v81, ClinVar 201706, NHLBI ESP V2-SSA137, HGMD-PUBLIC 20164, dbSNP150, GENCODE v27, gnomAD v2.1 and Regulatory Build. VEP provides scores and pathogenicity predictions with Sorting Intolerant From Tolerant v5.2.2 (SIFT) [28] and PolyPhen-2 v2.2.2 [29] tools. For each transcript in the final dataset we obtained the coding consequences prediction and score according to SIFT and PolyPhen-2. A canonical transcript was assigned for each gene, according to VEP.

### Serbian sample sex structure

To determine the sex structure of the Serbian population sample we used the CNVkit 0.9.1 toolkit [35]. We evaluated the number of mapped reads on sex chromosomes of each sample BAM file using the CNVkit to generate target and antitarget BED files.

### Description of Variants

In order to investigate allele frequency distribution in the Serbian population sample, we classified variants into four categories according to their minor allele frequency (MAF): MAF ≤ 1%, 1-2%, 2-5% and ≥ 5%. We classified separately singletons (MAC=1) and private doubletons (MAC=2), where a variant occurs only in one individual and in the homozygotic state.

We classified variants into four functional impact groups according to Ensembl (http://grch37.ensembl.org/info/genome/variation/prediction/predicted_data.html): HIGH (Loss of function) that includes splice donor variants, splice acceptor variants, stop gained, frameshift variants, stop lost and start lost. MODERATE that includes inframe insertion, inframe deletion, missense variants. LOW that includes splice region variants, synonymous variants, start and stop retained variants. MODIFIER that includes coding sequence variants, 5’UTR and 3’ UTR variants, non-coding transcript exon variants, intron variants, NMD transcript variants, non-coding transcript variants, upstream gene variants, downstream gene variants and intergenic variants.

In order to investigate the rate of overlap with reference databases and to determine novel variants, we compared our dataset with publicly available reference databases: gnomAD v3.0 genome (https://gnomad.broadinstitute.org/) [9], NHLBI Exome Sequencing Project database vesp6500 (https://esp.gs.washington.edu/drupal/) and the European population from the 1000 Genomes Project Phase 3 (1kGP_EUR) (https://www.internationalgenome.org/home) [7]. To compare our dataset with the previously mentioned databases, we used ANNOVAR v2019Oct24 [61] with a filter-based annotation option. Overlapping variants were identified comparing start and end positions, as well as the same observed alleles. Furthermore, we used BedTools v2.29.2 [62] to intersect multiple outputs from ANNOVAR to find variants that were not overlapping with any of the databases.

### Population Genetics Analyses

Principal Component Analysis (PCA) was used in order to estimate the ancestry of the Serbian population sample relative to other world populations. PCA was performed using the PLINK v1.9 (www.cog-genomics.org/plink/1.9/) [59] software on the Serbian population sample and 1000 Genomes Project Phase 3 (1kGP) [7] dataset as a reference dataset. 5473 autosomal SNVs in common between the Serbian population and 1kGP reference database were selected for the analysis.

To ensure high quality of variants, study data was pruned for variants that are in linkage disequilibrium (LD). The LD-pruned dataset was generated using the PLINK v1.9 --indep-pairwise option, with parameters of the 50kb window in which LD is calculated between each SNV pair and with the r^2^ > 1.5 threshold above which SNPs were removed. To ensure that only common (MAF≥ 5%) variants are considered in the analysis, we set --maf to 0.05. The number of SNVs to shift the window at each step was set to 5. To reduce the size of the reference dataset to the size of the Serbian population sample, we filtered the reference dataset with the Serbian population sample SNVs. In order to compute the joint principal components of the reference and study population, the two datasets were merged and PCA was performed on the combined data. We performed visualization of the PCA plot in Tableau v2020.1 (https://www.tableau.com/).

### Overrepresented Variants

To examine the variants that are significantly overrepresented in the Serbian population sample compared to European populations of 1kGP, we calculated the fold increase in AF as AF_(Serbian)_/AF_(European)_ and singled out variants with a fold increase > 5.

### Gene Ontology Annotations

For identification of biological processes in which the 16 genes with frequent variants in the Serbian population are involved, the Biological Process Ontology (BPO) in Gene Ontology (GO) was used, release 2020-07-16. GO terms were filtered by Evidence Codes: EXP, IDA, IPI, IMP, IGI, IEP, TAS and IC.

### Functional analysis of missense variants

Missense variants frequent in the Serbian population were analysed using the MutPred2 web server [37]. MutPred2 is a machine-learning method for predicting pathogenicity of amino acid substitutions (AAS), which integrates genetic and molecular data. It provides a general pathogenicity prediction, represented by the MutPred2 score, and a ranked list of specific molecular alterations potentially affecting the phenotype. MutPred2 estimates involvement of a missense variant in several structural and functional properties, including secondary structure, signal peptide and transmembrane topology, catalytic activity, macromolecular binding, post-translational modifications, metal-binding and allostery. We used the MutPred2 score ≥ 0.5 for predicting pathogenicity of a missense variant and p value ≤ 0.05 for predicting affected molecular mechanisms and motives.

Additional hypotheses about the functional effects of missense variants frequent in the Serbian population were created through a comprehensive manual search of available literature.

## Code Availability

We provided Common Workflow Language (CWL) [63] code workflow that was obtained and used on the Seven Bridges Cancer Genome Cloud platform in order to perform variant calling according to GATK v4.1 best practice recommendations, as Supplementary information json format files Supplementary_file_1.json, Supplementary_file_2.json and Supplementary_file_3.json.

Supplementary_file_1.json contains cwl workflow code that we used for data preprocessing and variant calling with Haplotype caller in ERC GVCF mode, Supplementary_file_2.json contains cwl workflow code for joint calling and Supplementary_file_3.json contains cwl workflow code that we used to perform hard filtering.

## DATA AVAILABILITY

The dataset generated and analysed during the current study is deposited and available as a single multisample VCF file at European Variation Archive (EVA), https://www.ebi.ac.uk/ena/browser/view/ERA3199532.

## AUTHORS’ CONTRIBUTION

N.V. designed the study. K.K.Lj. and B.Z. carried out sample sequencing. T.D. and V.K. processed data. T.D., V.P. and B.G. analysed data. V.P. was involved in development of software methods. B.G., M.L. and S.P. contributed to the interpretation of the results. T.D., B.G. and N.V. wrote the article. T.D. and V.P. prepared the figures. All authors have read and approved the submitted version.

