## Supplementary information for "The first insight into the genetic structure of the population of modern Serbia"

**for**

*Joint Corresponding authors

Average Coverage

Average coverage was calculated per position per sample, using SAMtools v.1.3. [1] and the overall population sample average coverage is 88.25. Supplementary Figure S1 shows the average coverage of each sample after target exome sequencing that is calculated per position and percent of coverage higher than the limit (20X) for each sample. Considering that the vast majority of samples show a large percent of coverage higher than limit, average coverage of samples indicates the appropriate quality of sequencing data.

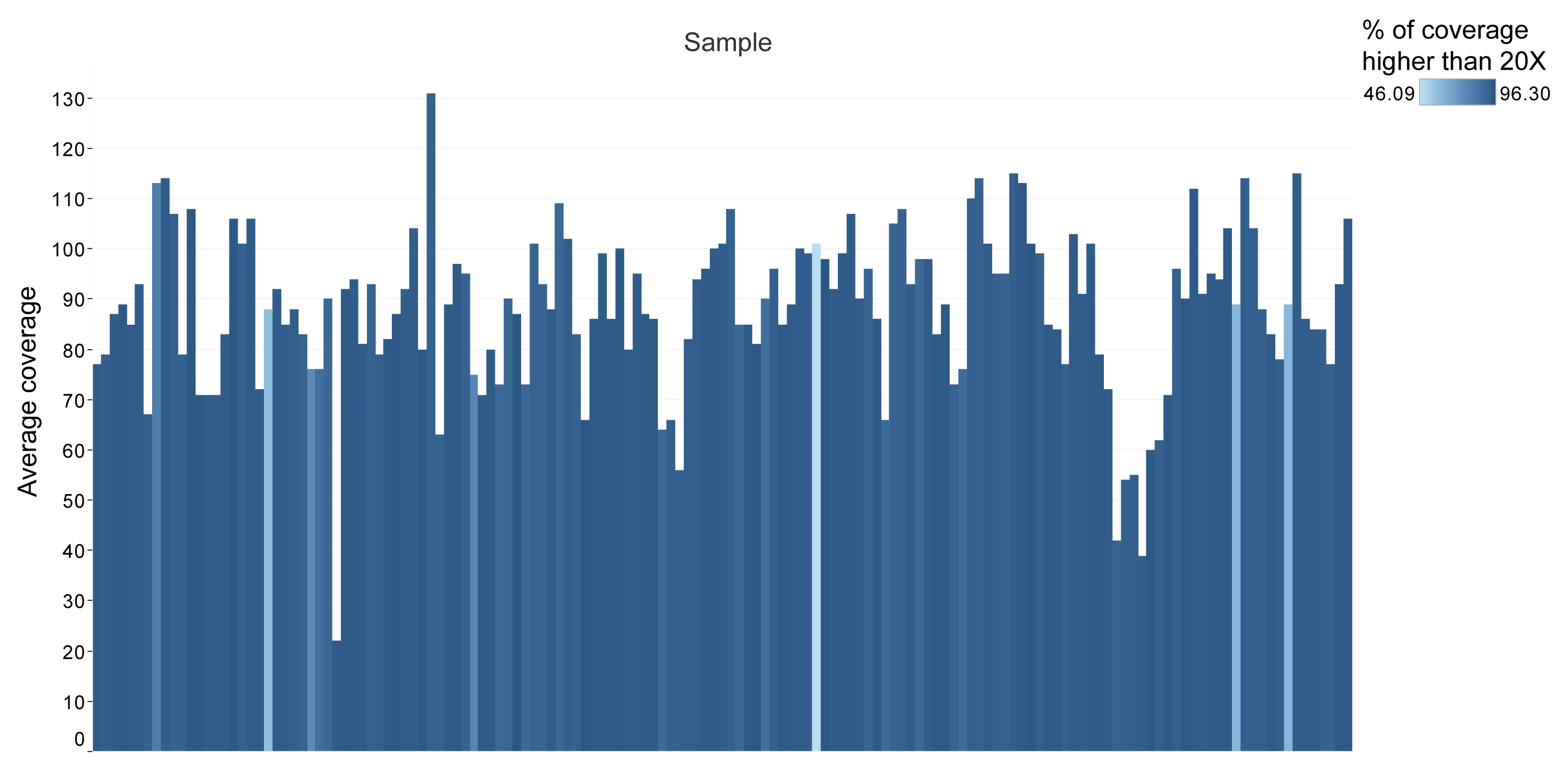

**Supplementary Figure S1** Target exome sequencing average coverage per sample. The y-axis represents the average coverage calculated per position for each sample and the x-axis with each bar represents individual samples. The color scale is used to represent percent of coverage higher than the limit (20X) for each sample. The percent of coverage higher than the limit for each sample in this study is in the range of 46.09 to 96.30.

Indel Distribution

A total of  1987 indels was discovered in the Serbian population sample, out of which 697 were in lengths divisible with 3, ranging from 3 to 60 bp. We marked indels that have lengths divisible by 3 as 3n indels and indels not divisible by 3 as non-3n indels, where n is an integer [2]. We expect that 3n indels do not cause frameshift mutation. Indel distribution shape (Supplementary Figure S2.) indicates an expected high number of 1 bp indels, considering the high frequency of short indels [3] and the possibility of PCR errors. According to Variant Effect Predictor (VEP) [4] annotation, in Serbian population sample 552 indels were described with frameshift variant consequence.

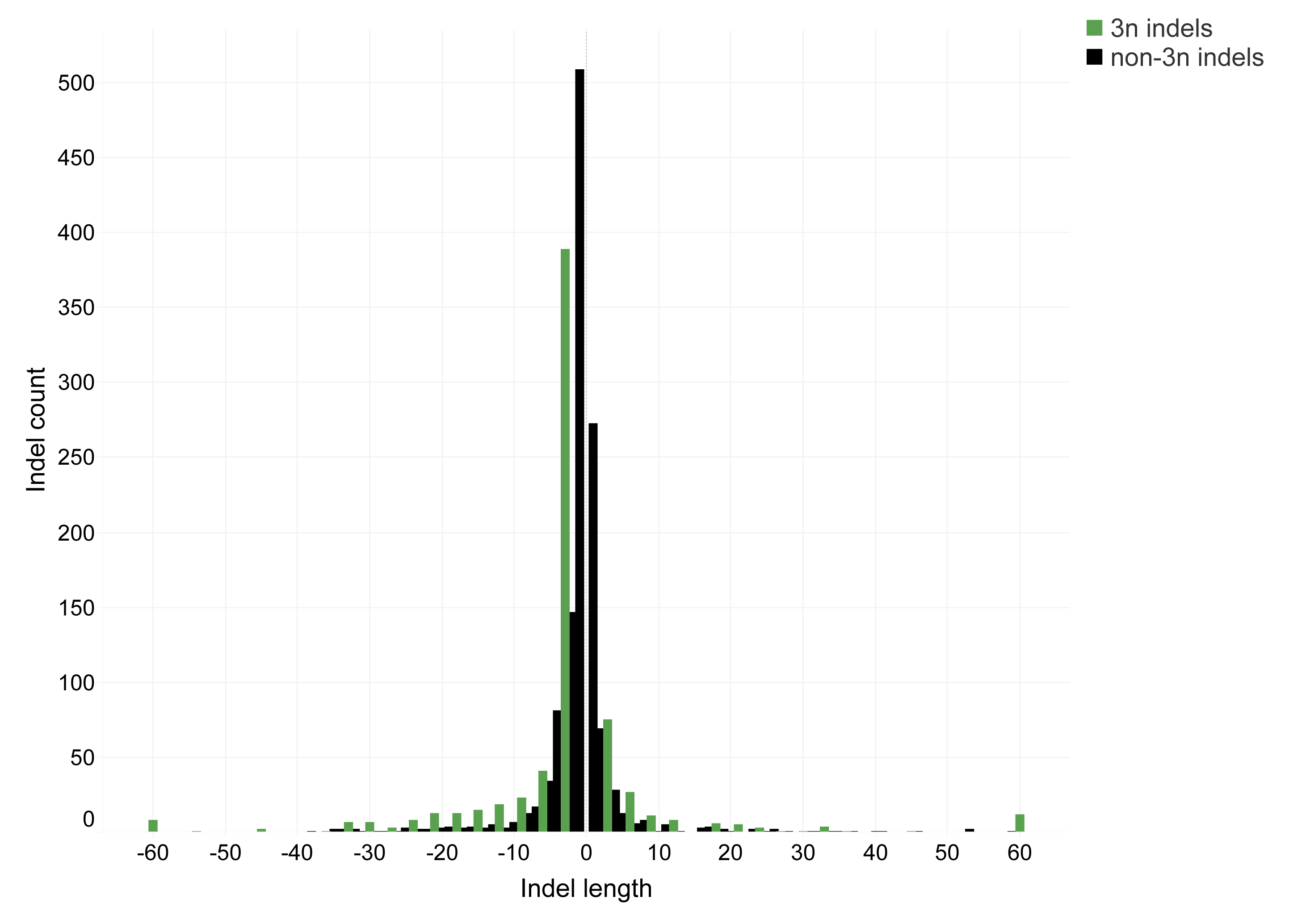

**Supplementary Figure S2.** Indel distribution. Indel length on the x-axis represents the length of each indel, thus Indel length > 0 shows insertions and Indel length < 0 shows deletions. Indel count on the y-axis shows the number of indels of a certain length. Shape of indel distribution demonstrates a high number of 1 bp indels, as well as indels with lengths divisible with 3, which are labeled with green as 3n indels. Indels not divisible by 3 are labeled as non-3n indels.

Pathogenicity prediction of variants using SIFT and PolyPhen-2 tools

The distribution of variants by SIFT [5] and PolyPhen-2 [6] prediction categories is shown in Supplementary Figure S3. SIFT tool predicts tolerated variants as most represented (Supplementary Fig. S3a) with 57.36%, additionally PolyPhen-2 tool supports this prediction with 66.67% benign variants (Supplementary Fig. S3b).

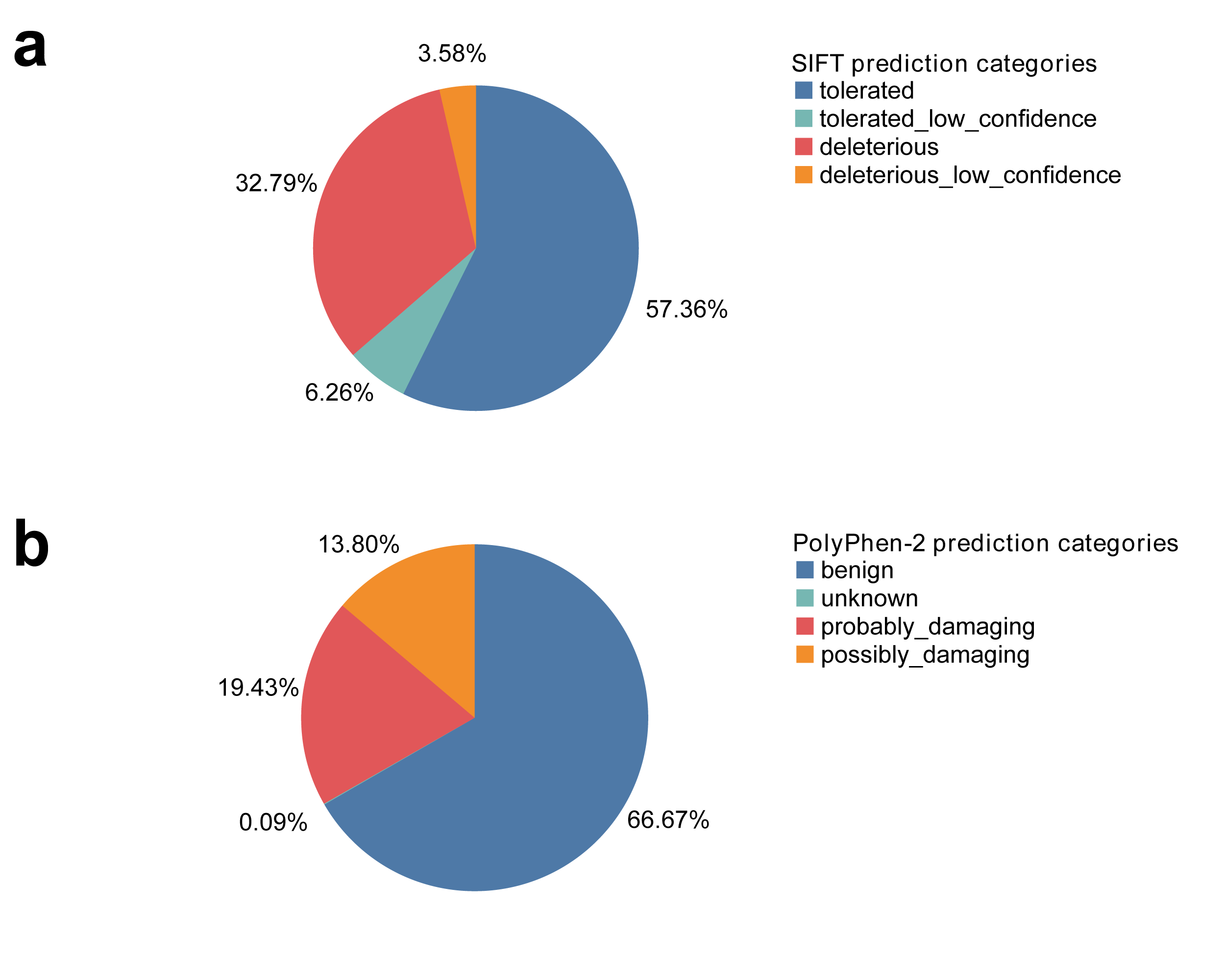

**Supplementary Figure S3** Distribution of variants by SIFT [5] and PolyPhen-2 [6] pathogenicity prediction categories. **a**. Distribution of variants by SIFT tool pathogenicity prediction categories with tolerated variants showed to be most abundant. **b.** Distribution of variants by PolyPhen-2 tool prediction categories, with benign variants prevailing.

**SUPPLEMENTARY TABLES**

| **Functional iImpact** | | | | | |
| --- | --- | --- | --- | --- | --- |
|  | HIGH | MODERATE | LOW | MODIFIER | **Total MAF** |
| Singletons | 690 (1.46) | 8911 (18.83) | 6439 (13.60) | 2541 (5.37) | 18581 (39.26) |
| Doubletons | 27 (0.06) | 241  (0.51) | 127 (0.27) | 91  (0.20) | 486 (1.03) |
| MAF<≤1% | 131 (0.27) | 2553  (5.40) | 2201 (4.65) | 766 (1.62) | 5651 (11.94) |
| MAF1-2% | 89 (0.18) | 2031  (4.30) | 1922 (4.06) | 730 (1.54) | 4772 (10.08) |
| MAF2-5% | 57 (0.12) | 1783  (3.76) | 1932 (4.08) | 700 (1.47) | 4472 (9.45) |
| MAF>≥5% | 99 (0.21) | 4399  (9.30) | 6713 (14.18) | 2151 (4.54) | 13362 (28.24) |
| **Total Impact** | 1093 (2.31) | 19918 (42.09) | 19334 (40.86) | 6979 (14.75) | 47324 (100) |

**Supplementary Table S1** The number of variants (and percentage in brackets) found in Serbian population sample, categorized by functional impact and Minor Allele frequency (MAF).

| **PolyPhen-2** | benign | Probably_  damaging | Possibly_  damaging | unknown |
| --- | --- | --- | --- | --- |
| **SIFT** |  |  |  |  |
| tolerated | 9409  (49.29%) | 621  (3.25%) | 915  (4.79%) | 5  (0.03%) |
| Tolerated_  low_  confidence | 1107  (5.80%) | 32  (0.17%) | 49  (0.26%) | 7  (0.04%) |
| deleterious | 1799  (9.42%) | 2923  (15.31%) | 1537  (8.05%) | 1  (0.005%) |
| Deleterious_  low_  confidence | 410  (2.15%) | 138  (0.72%) | 134  (0.70%) | 2  (0.01%) |

**Supplementary Table S2** The number of variants that overlap between SIFT [5] and PolyPhen-2 [6] functional prediction categories and percentage in brackets.

| **Gene** | **Variant** | **HGNC** | **Functional impact** | **Substitution type** | **% of variants in female sample** | **% of variants in male sample** | **Allele frequency** | | **Fold increase** |
| --- | --- | --- | --- | --- | --- | --- | --- | --- | --- |
|  |  |  |  |  |  |  | **1kGP** | **Serbian** |  |
| PSPH | rs79451216 | chr7:g.56019730G>A | MODERATE | missense | 36% | 30% | 0.001 | 0.163 | 163 |
| PCDHB4 | rs147934595 | chr5:g.141123572C>A | MODERATE | missense | 43% | 46% | 0.003 | 0.222 | 74 |
|  | rs144689137 | chr5:g.141123435G>A | LOW | synonymous | 38% | 45% | 0.008 | 0.208 | 26 |
| MRC1 | rs71497224 | chr10:g.17849727C>T | LOW | synonymous | 93% | 94% | 0.0109 | 0.771 | 70.73 |
|  | rs374113136 | chr10:g.17833763G>A | LOW | synonymous | 67% | 61% | 0.0497 | 0.41 | 8.25 |
| MYO15A | 17:181208 | chr17:g.18120828C>A | LOW | synonymous | 0 | 1% | 0.002 | 0.1 | 50 |
| KIR2DL1 | rs79002558 | chr19:g.54773524A>G | MODERATE | missense | 15% | 11% | 0.002 | 0.066 | 33 |
|  | 19:54775225 | chr19:g.54775225A>G | MODERATE | missense | 21% | 13% | 0.004 | 0.085 | 21.25 |
|  | 19:54775226 | chr19:g.54775226T>C | LOW | synonymous | 21% | 13% | 0.004 | 0.085 | 21.25 |
| KIR3DL1 | rs149760111 | chr19:g.54821568C>T | MODERATE | missense | 36% | 36% | 0.0149 | 0.198 | 13.29 |
| KIR2DL3 | rs149224427 | chr19:g.54742075C>G | MODERATE | missense | 54% | 52% | 0.0318 | 0.356 | 11.19 |
| HLA-DRB5 | rs147439581 | chr6:g.32518589C>T | MODERATE | missense | 57% | 65% | 0.0398 | 0.433 | 10.88 |
|  | rs41559420 | chr6:g.32519465G>A | MODERATE | missense | 20% | 22% | 0.0239 | 0.122 | 5.10 |
| CDSN | rs145583110 | chr6:g.31116168C>T | MODERATE | missense | 10% | 13% | 0.006 | 0.059 | 9.83 |
| RHPN2 | rs147870656 | chr19:g.32999658G>T | MODERATE | missense | 13% | 11% | 0.007 | 0.059 | 8.43 |
| BTNL2 | rs28362679 | chr6:g.32396116G>A | MODERATE | missense | 23% | 22% | 0.0159 | 0.115 | 7.23 |
| DND1 | 5:140671341 | chr5:g.140671341G>A | LOW | synonymous | 16% | 14% | 0.0109 | 0.077 | 7.06 |
| HTR3A | rs33940208 | chr11:g.113975355C>T | LOW | synonymous | 7% | 13% | 0.008 | 0.052 | 6.5 |
| GAL3ST3 | rs147282818 | chr11:g.66043629C>T | LOW | synonymous | 21% | 13% | 0.0139 | 0.09 | 6.47 |
| TNXB | rs17207895 | chr6:g.32052735T>C | MODERATE | missense | 11% | 25% | 0.0189 | 0.101 | 5.34 |
| HLA-DQB1 | rs41552812 | chr6:g.32664912C>T | MODERATE | missense | 21% | 23% | 0.0239 | 0.125 | 5.23 |

**Supplementary Table S3** Variants detected as frequent (MAF≥5) in Serbian population compared to European population of 1kGP and their sex representation.

| **Gene** | **Variant** | **AAS** | **MutPred2 score** | **MutPred2 - Affected PROSITE and ELM Motifs** | **MutPred2 - Molecular mechanisms** |
| --- | --- | --- | --- | --- | --- |
| PSPH | rs79451216 | R49W | 0.637 | - ELME000063 - CK1 Phosphorylation site - ELME000102 - NRD cleavage site - ELME000108 - PCSK cleavage site - PS00005 - Protein kinase C phosphorylation site | - Loss of Relative solvent accessibility - Loss of ADP-ribosylation at R49 - Altered Metal binding |
|  |  | R49G | 0.725 |  | - Loss of Relative solvent accessibility - Loss of Helix - Gain of ADP-ribosylation at R50 - Altered Metal binding, Gain of Methylation at R50 |
| KIR2DL1 | rs79002558 | S88R | 0.573 | - ELME000008 - PKA Phosphorylation site - ELME000012 - di Arginine retention/ retrieving signal - ELME000062 - PKA Phosphorylation site - ELME000063 - CK1 Phosphorylation site - ELME000070 - N-glycosylation site - ELME000102 - NRD cleavage site - ELME000108 - PCSK cleavage site | - Loss of Relative solvent accessibility - Altered Trans-membrane protein - Altered Ordered interface - Gain of Allosteric site at R89 - Gain of ADP-ribosylation at S88 - Altered DNA binding - Altered Metal binding - Loss of N-linked glycosylation at N84 |
| BTNL2 | rs28362679 | S334L | 0.514 | - ELME000053 - GSK3 phosphorylation site - ELME000062 - PKA Phosphorylation site - ELME000239 - USP7 binding motif - PS00007 - Tyrosine kinase phospho-rylation site 1 | - Altered Transmembrane protein - Loss of B-factor |
|  |  | S334W | 0.691 |  |  |
| HLA-DQB1 | rs41552812 | D89N | 0.503 |  | - Altered Ordered - Gain of Relative solvent accessibility |

**Supplementary Table S4** Variants detected as frequent in Serbian population that MutPred2 [7] predicted as affecting protein function with MutPred2 score and Affected PROSITE and ELM Motifs**.**

| **Gene** | **GO-BPO annotations** |
| --- | --- |
| PSPH | - L-serine metabolic process |
| PCDHB4 | - chemical synaptic transmission - nervous sustem development - synapse assembly |
| MRC1 | - Receptor-mediated endocytosis |
| KIR2DL1 | - natural killer cell inhibitory signalling pathway - regulation of immune response - immune response |
| KIR2DL3 | - immune response - regulation of immune response |
| KIR3DL1 | - natural killer cell mediated cytotoxicity - regulation of immune response |
| HLA-DRB5 | - antigen processing and presentation of exogenous peptide antigen via MHC class II - interferon-gamma-mediated signalling pathway - T cell receptor signalling pathway |
| DND1 | - negative regulation of gene silencing by miRNA - 3’UTR-mediated mRNA destabilization |
| CDSN | - corneocyte desquamation - skin morphogenesis cornification - cell adhesion - cell-cell adhesion - epidermis development - negative regulation of cornification |
| HTR3A | - chemical synaptic transmission |
| HLA-DQB1 | - Immunoglobulin production involved in immunoglobulin mediated immune response - humoral immune response mediated by circulating immunoglobulin - antigen processing and presentation of exogenous peptide antigen via MHC class II - T cell signalling pathway - interferon-gamma-mediated signalling pathway |
| TNXB | - collagen metabolic process - elastic fiber assembly |

**Supplementary Table S5** GO-BPO annotations for genes with variants overrepresented in the Serbian population sample. GO terms common for two or more genes are colored; red – immune response, blue – regulation of immune response, green – chemical synaptic transmission.
